# Identification of a novel breathing circuit that controls pain and anxiety

**DOI:** 10.1101/2020.01.09.900738

**Authors:** Shijia Liu, Mao Ye, Gerald M. Pao, Samuel Myeongsup Song, Jinho Jhang, Sung Han

## Abstract

Alleviating pain with controlled breathing has been practiced throughout human history. Despite its wide use and long history, a neural circuit-based understanding of the pain-breathing interaction is largely lacking. Here we report a novel breathing circuit that regulates non-homeostatic breathing rhythm, as well as pain and anxiety. We identify that a cluster of neurons expressing the *Oprm1* gene, which encodes the μ-opioid receptor (MOR) in the lateral subdivision of parabrachial nucleus (PBL^*Oprm1*^), directly regulates breathing rate in mice by conveying signals from the limbic areas to respiratory rhythm generating neurons in the medullary preBötzinger Complex (preBötC). In addition, we found that pain signals rapidly increase breathing rate by activating these neurons in both awake and anesthetized mice. Inactivating these neurons not only decreases the breathing rate, but it also substantially decreases anxiety-like behaviors and induces strong appetitive behaviors. Furthermore, PBL^*Oprm1*^ inactivation alleviates pain by attenuating the perception of the affective-motivational aspect of pain. These results suggest that PBL^*Oprm1*^ neurons play a critical role in the non-homeostatic regulation of breathing and in the regulation of pain and anxiety through breathing.

## Introduction

Breathing is a fundamental, life-sustaining function for animals^1^. Its rhythmicity is automatically generated and maintained homeostatically by rhythm-generating properties of a neuronal network in the medulla to sustain blood oxygen levels in response to changing oxygen needs^2^. However, breathing rhythms are also regulated by interoceptive and exteroceptive signals, such as emotional and sensory stimuli^3^. Pain is an emotional and sensory stimulus that greatly influences breathing behaviors. Intense and uncontrollable pain induces hyperventilation in humans^4^. Conversely, controlled breathing has been widely used as a non-pharmacological intervention for treating pain^5,6^, as well as affective disorders^5–8^. Although this reciprocal interaction between pain and breathing behaviors has been reported in a number of clinical and laboratory studies in human subjects^4^, investigations on molecular and circuit-based understanding of pain-breathing interactions is essentially lacking.

There are three respiration centers in the brain: the ventral respiratory group (VRG), the dorsal respiratory group (DRG), and the pontine respiratory group (PRG)^2^. The VRG and DRG are located in the medulla and act as generators of breathing rhythm and pattern^2,9,10^. However, the PRG, located in the pons, is not involved in generating respiratory rhythm, but has been proposed to have modulatory roles in response to interoceptive and exteroceptive signals^11^. This hypothesis is largely based on its anatomical location, which connects the forebrain and limbic structures to the medullary respiratory centers. However, supporting evidence for this hypothesis is limited. Although there are a few pioneering studies that have investigated non-homeostatic breathing behaviors^12,13^, the majority of research has focused on homeostatic regulation of respiratory networks. Therefore, to extend our knowledge for these respiratory behaviors, we should broaden the scope of research to interoceptive or exteroceptive regulation of breathing rhythm. Exploring the reciprocal interaction between pain and breathing will not only extend our knowledge of basic research on breathing behaviors, it will also broaden our understanding of breathing as a behavioral intervention of pain and anxiety. Here, we report that PBL^*Oprm1*^ neurons in the PRG play a key role in non-homeostatic breathing regulation and the pain-breathing interactions.

## Results

### Correlation of PBL^*Oprm1*^ activity and breathing rate

Whole-body plethysmography (WBP) is the most widely used tool for monitoring breathing behaviors^14^, because it provides multiple parameters for breathing behaviors with high precision. However, due to its closed configuration, it is difficult to access the brain during respiratory measurement. Moreover, its small chamber size hinders behavioral assessments for monitoring emotional behaviors in mice. To overcome this issue, we monitored breathing behaviors by measuring temperature fluctuation in the nasal cavity during gas exchange with a micro-thermistor implanted into the nasal cavities of test mice^15^. This approach may not provide as many breathing parameters as the WBP, but it renders unprecedented freedom to access the brain while simultaneously monitoring breathing behaviors. It also is possible to monitor breathing behaviors in unrestrained mice during behavioral tests. With this configuration, in combination with fiber photometry calcium imaging *in vivo*, we monitored activity of the PBL^*Oprm1*^ neurons and breathing behaviors in mice expressing a genetically encoded calcium indicator, jGCaMP7s, in their home cage (**Fig. 1A, B**). To express jGCaMP7s specifically in the PBL^*Oprm1*^ neurons, a recombinant adeno-associated virus that expresses jGCaMP7s (AAV-DIO-GCaMP7s) in a Cre-dependent manner was stereotaxically injected into the PBL of a Cre-driver mouse line that expresses the Cre recombinase in the *Oprm1*-expressing neurons (*Oprm1*^Cre/+^ mice) three weeks prior to the tests. Simultaneous measurements of neural activity and breathing behaviors showed a close correlation (**Fig. 1C, D, Fig. S1A-C**). Aligning the calcium traces with the onset of the breathing-rate increases show that single calcium events and breathing-rate increases are tightly correlated (**Fig. 1E**). Convergent cross-mapping (CCM) analysis, a nonlinear time series embedding algorithm that tests the causality of two independent time-series data^16^, can generate a model that predict the change in respiratory rate from calcium traces with greater than 95 % accuracy (**Fig. 1F, G**). These results suggest that PBL^*Oprm1*^ neurons are involved in gross changes of breathing rates.

**Figure 1.**
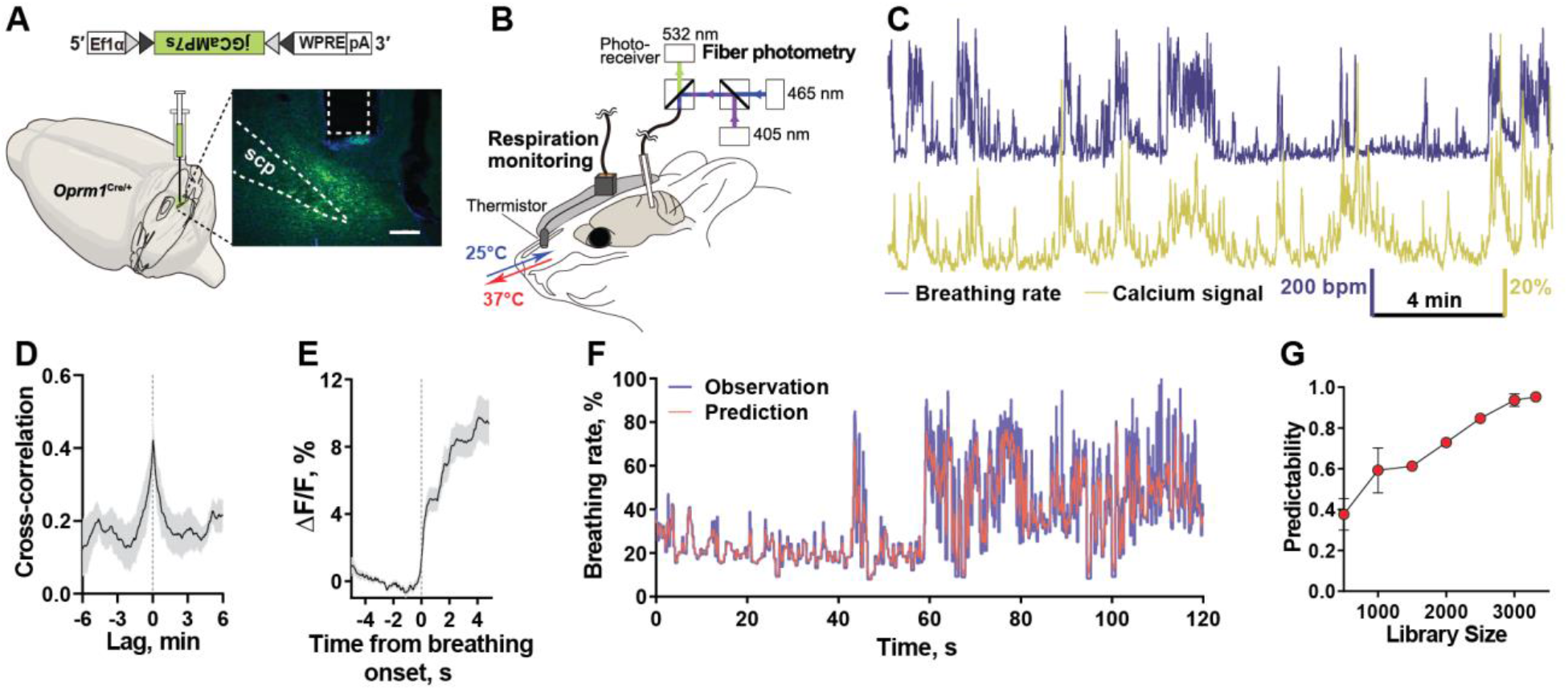
Tight correlation of PBL^*Oprm1*^ neuronal activity and breathing rates. **A**, Stereotaxic injection of AAV-DIO-jGCaMP7s into the PBL of the *Oprm1*^Cre/+^ mice. **B**, A schematic diagram of simultaneous monitoring of breathing rates and neural activity by thermistor sensor and fiber photometry. **C**, Breathing rates and calcium signals of PBL^*Oprm1*^ neurons are tightly correlated. **D**, Cross-correlation analysis of breathing rates and calcium signals (n = 4). **E**, Averaged trace of calcium events aligned to the onset of increased breathing rate. **F**, Convergent cross-mapping (CCM) analysis predicted the changes in breathing rate by the calcium traces with >90% accuracy. **G**, Predictability increased to > 90% as sample size increases (n = 4).

### PBL^*Oprm1*^ neurons regulate breathing rhythm

To test if changes in breathing rates are mediated by the PBL^*Oprm1*^ neurons, we manipulated these neurons selectively using both chemogenetic (with awake mice) and optogenetic approaches (with anesthetized mice). *Oprm1*^Cre/+^ mice were stereotaxically injected with either AAV-DIO-hM3Dq-mCherry (**Fig. S2A**) or AAV-DIO-hM4Di-mCherry (**Fig. S2E**) into the PBL to selectively activate or inhibit PBL^*Oprm1*^ neurons by injecting 5 mg/kg Clozapine-N-Oxide (CNO, 5 mg/kg), respectively. AAV-DIO-eYFP was injected as a control. Three weeks after injection, breathing rates were monitored with WBP in awake mice 30 minutes after CNO injection. Selective activation of PBL^*Oprm1*^ neurons in the hM3Dq-expressing group displayed significant increases in breathing rates as compared to the saline injected group, whereas no increase was observed in control eYFP-expressing mice (**Fig. S2B, C**). The action of CNO in this group was confirmed with *ex vivo* slice electrophysiological recordings with cell-attached configuration. Bath application of 3 μM CNO dramatically increased the firing rate of the PBL^*Oprm1*^ neurons in the hM3Dq-expressing *Oprm1*^Cre/+^ mice (**Fig. S2D**). Selective inactivation in the hM4Di-expressing group by CNO displayed a significant decrease in breathing rate compared to the saline injected group, whereas no decrease was observed in the control eYFP group. (**Fig. S2F, G**). Bath application of 3 μM CNO in the slice of the same group of mice showed complete silencing of action potential firing in whole-cell patch clamp configuration (**Fig. S2H**). These data indicate that PBL^*Oprm1*^ neurons are necessary and sufficient for modulating breathing rates in awake, behaving mice. We wondered whether these neurons can also regulate breathing rates in lightly anesthetized mice. We used optogenetics to test this idea. We first injected AAV-DIO-ChR2-eYFP into the PBL of the *Oprm1*^Cre/+^ mice to selectively activate these neurons (**Fig. 2A**). We then monitored breathing rates in anesthetized mice with inductance plethysmography by placing piezo-based pressure sensors beneath the chests of test mice to detect chest expansion during breathing^17^. Optogenetic stimulation of these neurons significantly increased the breathing rates of the anesthetized mice (**Fig. 2B, C**). Optogenetic inhibition with ArchT (**Fig. 2D**) resulted in a significant suppression of breathing rates compared to the eYFP-expressing control group (**Fig. 2E, F**). Note that the basal breathing rate under anesthesia is much lower than in awake mice. These data suggest that PBL^*Oprm1*^ neurons control breathing behaviors, regardless of the conscious state of the test mice.

**Figure 2.**
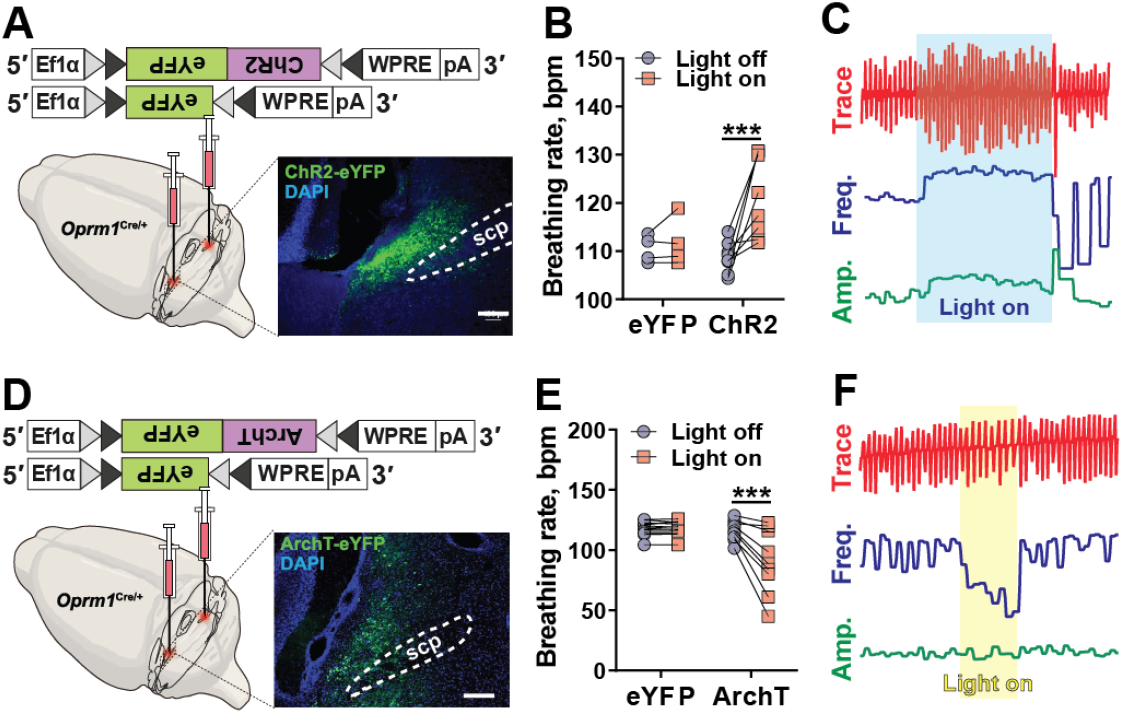
PBL^*Oprm1*^ neurons directly modulate breathing rates. **A**, Stereotaxic injection of AAV-DIO-ChR2-eYFP and AAV-DIO-eYFP into the PBL of the *Oprm1*^*Cre*/+^ mice. **B**, Inductance plethysmography showed that optogenetic stimulation of PBL^*Oprm1*^ neurons significantly increased breathing rates in ChR2-expressing anesthetized mice (n = 8), whereas eYFP-expressing control mice displayed no changes in breathing rates by light stimulation (n = 4). Two-way ANOVA analysis with Bonferroni’s multiple comparison post-hoc test, ***, *p* < 0.001. **C**, Upper panel, representative voltage trace representing breathing behaviors for 20 s. Middle panel, breathing frequency difference calculated from the upper panel. Range: 52.36-201.89 rpm. Lower panel, breathing amplitude difference calculated from the upper panel. Range: 4.29-12.67 mV. Blue area indicates 10-s optogenetic stimulation (470 nm). **D**, Stereotaxic injection of AAV-DIO-ArchT-eYFP and AAV-DIO-eYFP into the PBL of the *Oprm1*^Cre/+^ mice. **E**, Inductance plethysmography showed that optogenetic inhibition of PBL^*Oprm1*^ neurons significantly decreased breathing rates in ArchT-expressing anesthetized mice (n = 13 stimulations from 3 mice), whereas eYFP-expressing control mice displayed no changes in breathing rates by light stimulation (n = 9 stimulations from 3 mice). Two-way ANOVA analysis with Bonferroni’s multiple comparison post-hoc test, ***, *p* < 0.001. **F**, upper panel, voltage trace representing breathing behaviors for 10 s. Middle panel, breathing frequency change calculated from the upper panel. Range 84.47-122.59 rpm. Lower panel, breathing amplitude change calculated from the upper panel. Range 6.63-8.60 mV, Yellow area indicates 5-s optogenetic stimulation (589 nm). Scale bar, 200 μm. Data are presented as mean ± SEM.

### Input/output pathways of PBL^*Oprm1*^ neurons

We investigated the brain regions that are connected with PBL^*Oprm1*^ neurons. To identify direct upstream neurons, we performed monosynaptic, retrograde tracing with G-deleted pseudotyped rabies virus (RVdG) (**Fig. 3A**)^18^. Recombinant AAV expressing avian sarcoma leucosis virus glycoprotein EnvA and rabies glycoprotein (AAV8-hSyn-FLEX-TVA-P2A-GFP-2A-oG) in a Cre-dependent manner were injected into the PBL of the *Oprm1*^Cre/+^ mice. Three weeks after the injection, RVdG encoding mCherry (RVdG-mCherry) was injected into the same area. These mice were euthanized and perfused seven days after the RVdG injection to identify brain areas that express mCherry. The results showed that there are multiple brain areas that are directed connected to PBL^*Oprm1*^ neurons (**Fig. 3B**), including limbic areas such as central nucleus of the amygdala (CeA), the bed nucleus of the stria terminalis (BNST), the parvocellular subparafascicular nucleus (SPFp), the parasubthalamic nucleus (PSTN), as well as midbrain/brainstem areas such as the nucleus tractus solitarius (NTS), the superior colliculus (SC) (**Fig. 3A**). Interestingly, these areas are known to either relay aversive sensory signals or integrate sensory information with emotional valence^19–22^. We also investigated downstream brain areas that receive direct input from the PBL^*Oprm1*^ neurons by stereotaxic injection of recombinant AAV that express ChR2:eYFP in a Cre-dependent manner (AAV-DIO-ChR2:eYFP), which is expressed in the cell body and axon terminals (**Fig. S3**). YFP fluorescence was observed in the CeA, BNST, SPFp, PSTN, and preBötC. Interestingly, most of these regions are reciprocally connected to and from the PBL^*Oprm1*^ neurons. Among these areas, preBötC is well-known as a central rhythm generator of breathing rhythm^23^. To confirm the axonal projections of these neurons to the preBötC, we stereotaxically injected recombinant AAV expressing nuclear tdTomato in a Cre-dependent manner and synaptophysin-eGFP fusion protein (AAV-DIO-tdT-Syn:eGFP), which is targeted to the axon terminals (**Fig. 3C**). We confirmed that the eGFP signals from axonal terminals of PBL^*Oprm1*^ neurons are colocalized with the somatostatin (SST)-positive signals, which are markers of preBötC neurons (**Fig. 3D**). We reasoned that the PBL^*Oprm1*^ neurons may regulate breathing rates by modulating the activity of rhythm generating neurons located in the preBötC. To test this hypothesis, we expressed hM3Dq into the PBL^*Oprm1*^ neurons by stereotaxically injecting AAV-DIO-hM3Dq-mCherry into the PBL of the *Oprm1*^Cre/+^ mice. Three weeks after injection, the axon terminals of PBL^*Oprm1*^ neurons in the preBötC area were activated by stereotaxic injection of 200 nl of clozapine (1 μg/ml)^24^ into the preBötC area (**Fig. 3F**). We included controls to ensure that clozapine selectively actives DREADD-expressing neurons. Selective stimulation of preBötC-projecting terminals from the PBL^*Oprm1*^ neurons resulted in a significant increase in breathing rates in anesthetized *Oprm1*^Cre/+^ mice (**Fig. 3E, G**). Collectively, these results suggest that the PBL^*Oprm1*^ neurons can regulate breathing behaviors by conveying signals from the limbic and midbrain / brainstem areas to preBötC rhythm generating neurons.

**Figure 3.**
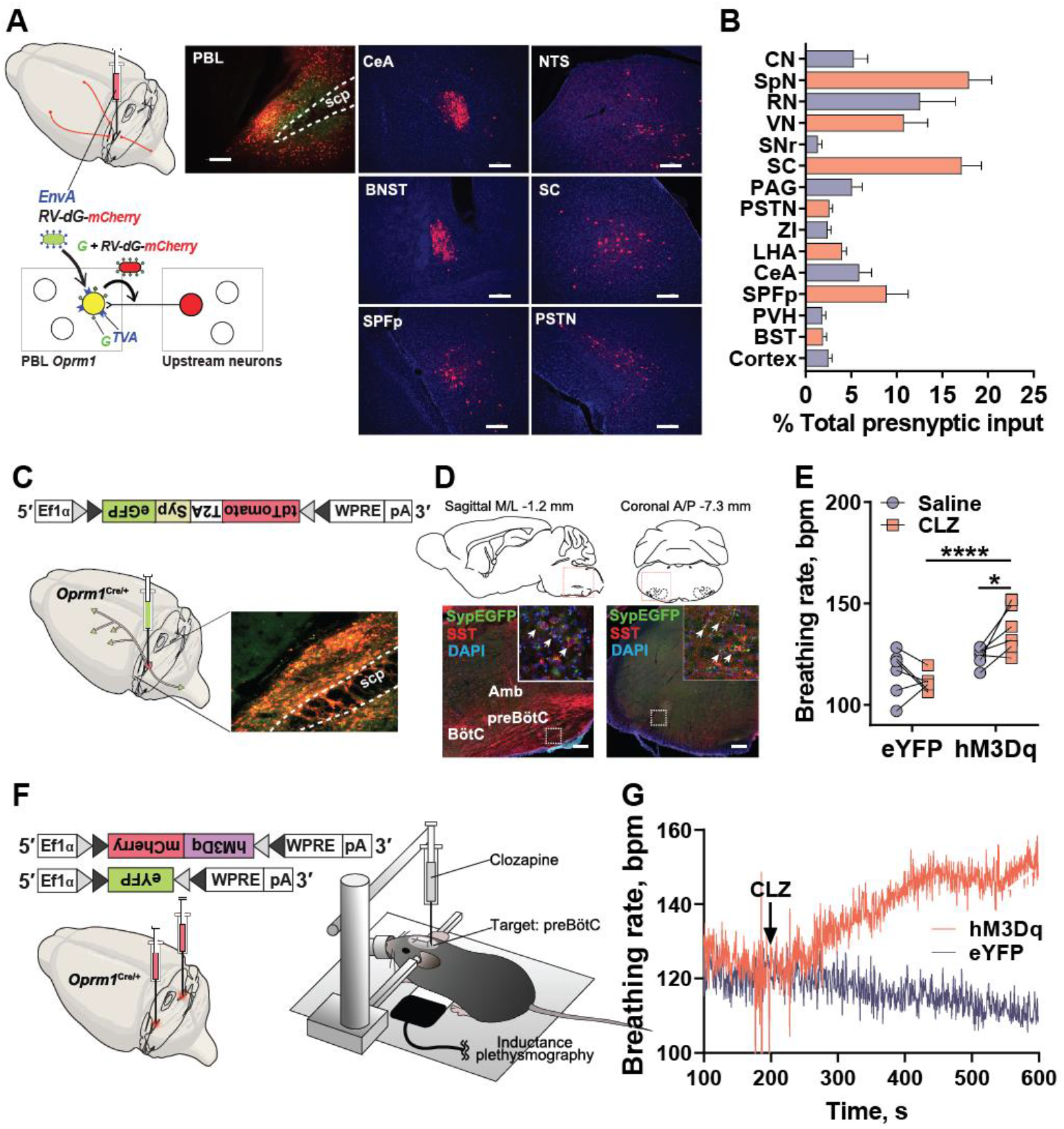
Input and output mapping of PBL^*Oprm1*^ neuronal connections. **A**, Monosynaptic retrograde tracing from PBL^*Oprm1*^ neurons using RVdG. Starter region (PBL) and example direct upstream areas with RVdG-mCherry expression. **B**, Quantification of direct upstream areas (n = 7). **C**, Illustration of output mapping by synaptophysin-eGFP expression in PBL^*Oprm1*^ neurons. **D**, PBL^*Oprm1*^ neurons make direct anatomical connection with the SST-positive preBötC neurons. **E**, Illustration showing chemogenetic stimulation of axonal terminal of hM3Dq-expressing PBL^*Oprm1*^ neurons though stereotaxic injection of 200 nl, clozapine (CLZ, 1 μg/ml) into the preBötC area. **F,** changes in breathing rate before and after clozapine injection. **G**, Chemogenetic stimulation of axonal terminals of hM3Dq-expressing PBL^*Oprm1*^ neurons increased breathing rate (n = 7) while eYFP-expressing group was unaltered (n = 6). Two-way ANOVA analysis with Bonferroni’s multiple comparison post-hoc test, *, *p* < 0.05. ****, *p* < 0.0001. Scale bar, 200 μm. Abbreviations: **A**, CeA: central amygdalar nucleus; BST: bed nuclei of the stria terminalis; SPFp: subparafascicular nucleus, parvicellular part; NTS: nucleus tractus solitarius; SC: superior colliculus; PSTN: parasubthalamic nucleus. **B**, CN: cochlear nucleus; SpN: spinal nucleus; RN: reticular nucleus; VN: vestibular nucleus; SNr: substantia nigra, reticular part; PAG: periaqueductal gray; ZI: zona incerta; LHA: lateral hypothalamic area; PVH: paraventricular hypothalamic nucleus.

### Pain increases breathing rates via PBL^*Oprm1*^ neurons

PBL^*Oprm1*^ neurons receive monosynaptic inputs from the brain areas that mediates pain perception and emotional regulation. Therefore, we tested if they are responsible for the reciprocal interaction of pain and breathing. We monitored breathing behavior and PBL^*Oprm1*^ activity in response to noxious sensory stimuli in anesthetized mice. Breathing rates were monitored with inductance plethysmography using a piezo sensor under the chests of anesthetized mice and calcium signals were monitored with fiber photometry (**Fig. 4A, F**). We stimulated the mice with thermal pain by touching a temperature-controlled rod (55 °C) to the tail of anesthetized mice (**Fig. 4A**). Thermal pain robustly increased both the breathing rates (**Fig. 4B, C**) and calcium signals (**Fig. 4D, E**). The same group of mice were also tested with mechanically noxious stimuli (Tail pinch with 300 g pressure) (**Fig. 4F**), which also robustly increased both breathing rates (**Fig. 4G, H**) and calcium signals (**Fig. 4I, J**). We also tested this hypothesis in freely moving mice using thermistor-based plethysmography in combination with fiber photometry in thermistor-implanted *Oprm1*^Cre/+^ mice that express jGCaMP7s in the PBL^*Oprm1*^ neurons (**Fig. S4A**). The test mice were placed on the surface of the hot plate and habituated at room temperature. Increasing the surface temperature from room temperature to 55°C induced dramatic increases in both breathing rates and calcium signals in awake mice (**Fig. S4B-D**). Cross-correlation analysis showed that these activities were tightly correlated (**Fig. S4E**). We then inhibited these neurons through optogenetic silencing to test whether these neurons are responsible for the increase in breathing rates by pain signals. The anesthetized *Oprm1*^Cre/+^ mice expressing ArchT in the PBL^*Oprm1*^ neurons were placed on the piezo sensor, and the PBL^*Oprm1*^ neurons were optogenetically silenced during the thermal- and mechanical-pain induction (**Fig. 4K**). Optogenetic inhibition of these neurons significantly decreased both the thermal (**Fig. 4L**) and mechanical (**Fig. 4M**) pain-induced increases in breathing rates. In contrast, it failed to decrease pain-induced breathing rate increase in the eYFP control group. These results show that the PBL^*Oprm1*^ neurons can mediate the increase in pain-induced breathing rate.

**Figure 4.**
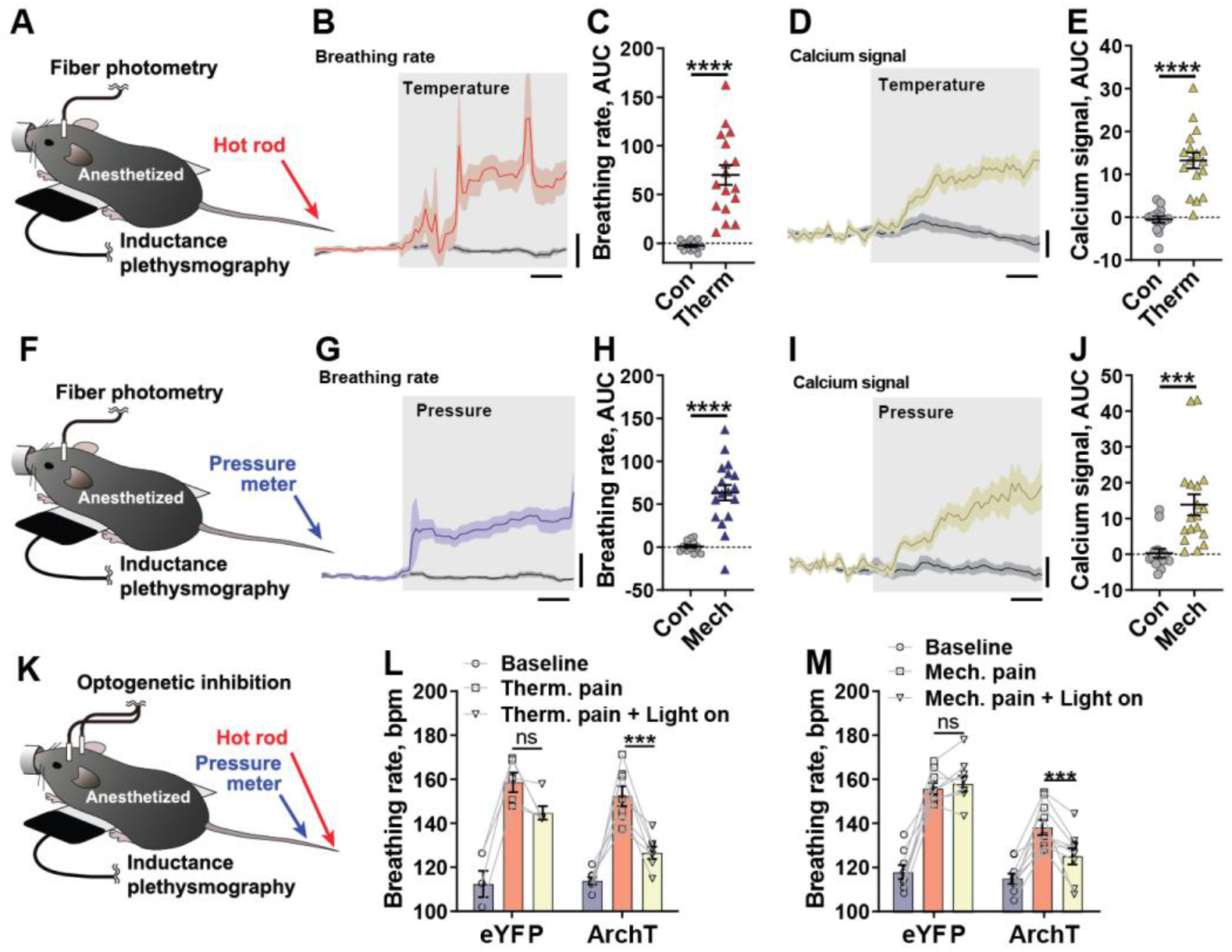
PBL^*Oprm1*^ neurons mediate pain-induced breathing rate increase. **A**, Simultaneous monitoring of PBL^*Oprm1*^ neuronal activity and breathing rates when anesthetized mice were stimulated with thermal pain (55°C hot rod). **B**, Average breathing-rate trace stimulated by hot rod (red) and room-temperature rod (grey). Scale bar, 1 s/10 bpm. **C**, Area Under Curve (AUC) analysis of the breathing-rate traces (n = 7, 3 repeated stimulation in each mouse), unpaired t-test, ****, *p* < 0.0001. **D**, Average calcium trace stimulated by hot rod and room-temperature rod. Scale bar, 1 s/2 % ΔF/F. **E**, AUC analysis of the calcium traces (n = 7, 3 repeated stimulation in each mouse), unpaired t-test, ****, *p* < 0.0001. **F**, Simultaneous monitoring activity of PBL^*Oprm1*^ neurons and breathing rate when anesthetized mice were stimulated with mechanical pain signal. **G**, Average breathing-rate traces stimulated by tail pinch with 300 g (blue) and 0 g (grey) pressure. Scale bar, 1 s/10 bpm. **H**, AUC analysis of the breathing rate traces (n = 7, 3 repeated stimulation in each mouse), unpaired t-test, ****, *p* < 0.0001. **I**, Average calcium trace stimulated by tail pinch with 300 g and 0 g pressure. Scale bar, 1 s/2 % ΔF/F. **J**, AUC of the calcium traces (n = 7, 3 repeated stimulation in each mouse), unpaired t-test, ***, *p* < 0.001. **K**, Monitoring breathing rate during optogenetic inhibition of PBL^*Oprm1*^ neurons by ArchT in anesthetized mice stimulated with thermal and mechanical pain signals. **L-M**, Optogenetic inhibition of PBL^*Oprm1*^ neurons significantly suppressed breathing rates evoked by thermal (**L**) and mechanical pain signals (**M**) (n = 13 stimulations from 3 mice), but remained unaltered in the eYFP-expressing control group (n = 9 stimulations from 3 mice), Two-way ANOVA analysis with Bonferroni’s multiple comparison post-hoc test. ns, not significant; ***, *p* < 0.001. Data are presented as mean ± SEM.

### Inhibiting PBL^*Oprm1*^ activity alleviates pain

Human studies have shown that slow breathing alleviates pain^5^. To test if decreases in breathing rates attenuate pain perception, we chemogenetically inhibited PBL^*Oprm1*^ neurons and performed somatosensory perception tests. To test thermal sensitivity, *Oprm1*^Cre/+^ mice expressing hM4Di in the PBL^*Oprm1*^ neurons were injected with CNO (7.5mg/kg) 45-60 min prior to the hot-plate test. Latency of paw withdrawal or licking on the hot plate was monitored. Latency was not significantly different compared to neurons expressing eYFP (**Fig. S5A, B**). Likewise, mechanical sensitivity with the same group of mice in automatic von Frey test showed no difference compared to eYFP-control mice (**Fig. S5C, D**). However, in the formalin test that evaluates chemical and inflammatory pain perception caused by continuous pain from injured tissue^25^ (**Fig. 5A**), hM4Di-expressing mice injected with CNO displayed significantly attenuated paw-licking behaviors. During initial 5-min after formalin injection, CNO-injected hM4Di-expressing mice display behavioral trend toward decreased licking (**Fig. 5B**). During inflammatory pain phase (20-35 min after formalin injection), the paw-licking behavior was significantly decreased in CNO-injected hM4Di-expressing mice when compared to control eYFP expressing mice (**Fig. 5C**). Unlike thermal and mechanical sensitivity tests, inflammatory pain monitored by formalin-induced paw licking behavior is associated with an affective and motivational component of pain^26^. Therefore, these results suggest that inactivating PBL^*Oprm1*^ neurons attenuates the affective-motivational aspect of pain without altering the sensory perception of pain. To further test this idea, we performed a fear-conditioning test (**Fig. 5D**). In classical Pavlovian, fear conditioning, foot-shock pain is a motivational signal to remember noxious cues or contexts. Therefore, freezing behavior after conditioning is directly correlated with perception of the affective-motivational aspect of pain. During the training session of contextual fear conditioning, the freezing behaviors of the control group of mice gradually increased as the 0.2-mA foot shock was repeatedly delivered, but the hM4Di-expressing mice displayed significant attenuation of freezing in response to the same stimuli (**Fig. 5E**). The attenuated freezing response of the hM4Di group persisted during the fear-memory test 24 hr after the training compared to eYFP control mice (**Fig. 5F**). An elevated-plus-maze (EPM) test showed that mice with chemogenetic inactivation of the PBL^*Oprm1*^ neurons spent significantly longer time (**Fig. 5G-I**) and visited open arms more frequently (**Fig. S5E, F**) compared with the control group. We also performed real-time place preference (RTPP) (**Fig. 5J**) and real-time place aversion (RTPA) (**Fig. 5M**) test with the mice expressing ArchT, and ChR2 in the PBL^*Oprm1*^ neurons to test whether inactivation, or activation of the PBL^*Oprm1*^ neurons encode positive, or negative valence, respectively. The ArchT-expressing mice stayed significantly longer time in the “light-on” chamber as compared with the “light-off chamber”, indicating that inactivating the PBL^*Oprm1*^ neurons is preferred, whereas the control mice expressing eYFP spent equal time in both chambers (**Fig. 5K, L**). In contrary, the ChR2-expressing mice displayed avoidance behavior towards the “light-on” chamber (**Fig. 5N, O**). Overall, these results indicate that PBL^*Oprm1*^ inactivation alleviates the perception of the affective-motivational aspect of pain, leaving the perception of sensory-discriminative pain unaltered by encoding positive valence in mice.

**Figure 5.**
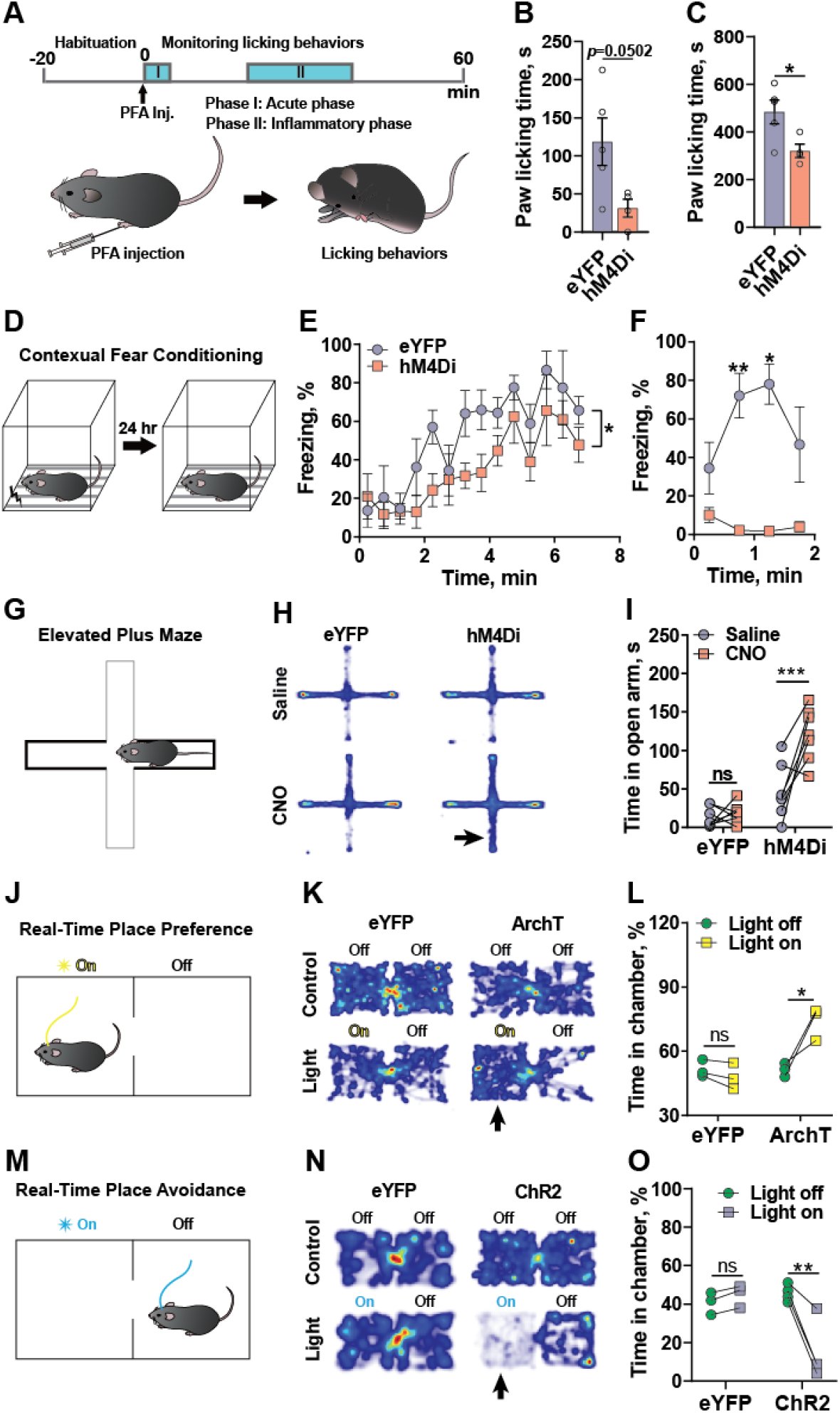
Inhibition of PBL^*Oprm1*^ neuronal activity alleviates affective-motivational pain perception and anxiety-like behaviors. **A**, Formalin assay for chemical and inflammatory sensitivity test after CNO injection in mice expressing eYFP or hM4Di in PBL^*Oprm1*^ neurons (n = 5). **B**, Chemical pain sensitivity test by monitoring paw licking behavior during initial 5 minutes after formalin injection on the plantar surface of a hind paw. Unpaired t-test, *p* = 0.0502. **C**, Inflammatory pain sensitivity test by monitoring paw licking behavior 20-35 min after formalin injection. Unpaired t-test, *, *p* < 0.05. **D**, Contextual fear-conditioning test after CNO injection prior to conditioning in mice expressing eYFP or hM4Di in PBL^*Oprm1*^ neurons. **E**, Percent freezing behaviors during conditioning by 5 consecutive foot shocks (0.2 mA, 2 s) with 1-minute inter-shock interval (n = 5), Two-way ANOVA analysis with Bonferroni’s multiple comparison post-hoc test. *, *p* < 0.05. **F**, Percent freezing behavior during contextual test 24 hr after the conditioning (n = 5). Two-way ANOVA analysis with Bonferroni’s multiple comparison post-hoc test, *, *p* < 0.05; **, *p* < 0.01. **G**, Elevated-plus-maze (EPM) test with mice expressing hM4Di or eYFP in PBL^*Oprm1*^ neurons. **H**, Heat maps representing average movement of test mice in each group during EPM test. **I**, Mice expressing hM4Di stayed significantly longer time in the open arm when injected with CNO (7.5 mg/kg) compared to saline injection, whereas the eYFP-expressing control group showed no significant increase of time in the open arm by CNO or saline injection (n = 7). Two-way ANOVA analysis with Bonferroni’s multiple comparison post-hoc test. ns, not significant; ***, *p* < 0.001. **J**, Real-time place preference (RTPP) test with mice expressing ArchT or eYFP in PBL^*Oprm1*^ neurons. **K**, Heat maps representing average movement of test mice in each group during RTPP test. **L**, Mice expressing ArchT displayed significant preference to the “light-on” chamber compared to the “light-off” chamber, whereas the eYFP-expressing control group showed no preference (n = 3), Two-way ANOVA analysis with Bonferroni’s multiple comparison post-hoc test. ns, not significant. *, *p* < 0.05. **M**, Real-time place avoidance (RTPA) test with mice expressing ChR2 or eYFP in PBL^*Oprm1*^ neurons. **N**, Heat maps representing average movement of test mice in each group during RTPA test. **O**, Mice expressing ChR2 (n = 4) displayed significant avoidance to the “light-on” chamber (40 Hz) compared to the “light-off” chamber, whereas the eYFP-expressing control group showed no preference (n = 3), Two-way ANOVA analysis with Bonferroni’s multiple comparison post-hoc test. ns, not significant. **, *p* < 0.01.

## Discussion

We identified a genetically defined circuit that directly regulates breathing rhythm by conveying signals from the limbic / brainstem brain areas to the medullary rhythm generating neurons. Activity of PBL^*Oprm1*^ neurons are tightly correlated with breathing-rate changes in awake, behaving mice, and manipulating the activity of these neurons directly regulates breathing rates in both awake and anesthetized mice. There is a direct connection from PBL^*Oprm1*^ neurons to the preBötC that is responsible for breathing regulation. Furthermore, we show that pain signals increase breathing rates through this ponto-medullary circuit. Suppressing the activity of this circuit alleviates the affective-motivational aspect of pain, as well as anxiety-like behaviors. Therefore, these findings suggest that inhibiting the PBL^*Oprm1*^ to preBötC pathway may be one of the mechanisms by which controlled breathing suppresses pain.

Notably, pain-induced increases in breathing rate occurred in anesthetized mice, and this increase was blocked by inhibiting the PBL^*Oprm1*^ neurons. These results suggest that the pain-to-breathing modulation is an autonomic response. Conversely, it is unclear whether breathing-to-pain modulation is an autonomic or non-autonomic response. Decreased chest movement itself may involuntarily generate a sensory feedback that modulates the cognitive perception of pain. Alternatively, the brain’s voluntary command to decrease breathing rates directly regulates the activity of PBL^*Oprm1*^ neurons, thereby simultaneously generating physiological and emotional outputs. To address this question, it will be necessary to develop a behavioral or physiological paradigm that changes breathing rhythms involuntarily, like the Iron-lung for polio patients.

The identification of the neural circuit that simultaneously regulates both the emotional response of pain and breathing behaviors not only provides a circuit-based understanding of reciprocal breathing and pain modulation, but it also suggests that emotion and physiology may be simultaneously regulated through a common upstream circuit^27,28^.

## Materials and methods

### Animals

All protocols for animal experiments were approved by the IACUC of the Salk Institute for Biological Studies according to NIH guidelines for animal experimentation. The *Oprm1-Cre:GFP* transgenic mouse line used in this study was generated from the lab of Dr. Richard Palmiter^1^. Both male and female mice were used in all studies. Animals were randomized to experimental groups and no sex differences were noted. Mice were maintained on a 12 h light/dark cycle and provided with food and water *ad libitum*.

### Respiratory measurements

#### Inductance plethysmography

Inductance plethysmography (Fig 1k-l, p-q, Fig 2e-g, Fig 3a-j, p-r) was performed by placing a piezoelectric film beneath the chest of an anesthetized animal, which converts the chest movements into voltage signals. The PowerLab system with LabChart 8 software (ADInstruments Inc., USA) was used for data acquisition, inspiratory and expiratory peak detection and rate and amplitude calculation. Data were sampled at 400 Hz, low-pass filtered at 10 Hz, and smoothed with a 100-ms moving window. Automatic peak detection was validated with manual peak detection.

#### Whole body plethysmography (WBP)

A custom-built WBP chamber was utilized for measuring respiratory changes (Fig 1i, n). The PowerLab system with LabChart 8 software was used for data acquisition, inspiratory and expiratory peak detection and rate and amplitude calculation. Data were sampled at 1-kHz, band-pass filtered at 1-10 Hz, and smoothed with a 100-ms moving window. Automatic peak detection was validated with manual peak detection.

Mice were introduced into the WBP chamber for three 20 min habituation sessions before testing. Mice were kept in the chamber for 10-12 min during the testing session before and after the drug injection. After 5-10 min of chamber introduction, a stable pattern was reached and the averaged value of a stabilized 1-min segment was analyzed.

#### Micro thermistor-based plethysmography

A custom-built micro thermistor (Fig 1b-g, Fig 3k-o) was implanted into the mouse nasal cavity to detect changes in temperature between inspiratory and expiratory airflow^2^. The sensor was assembled using a Negative Temperature Coefficient (NTC) thermistor (TE Connectivity Ltd., Switzerland), an interconnector (Mill-Max Mfg. Corp., USA), and a voltage divider (Phidgets Inc., Canada). PowerLab (ADInstruments Inc., USA) was used for data acquisition, inspiratory and expiratory peak detection and rate and amplitude calculation. Data were sampled at 1 kHz, filtered with a 0.4-25 Hz band-pass filter, and smoothed with a 50 ms moving window. Automatic peak detection was validated with manual peak detection.

### Stereotaxic surgery

Mice were anesthetized with isoflurane (5% induction, 1.5-2% maintenance with a nose cone; Dräger Vapor 2000, Draegar, Inc., USA) and placed onto a water recirculating heating pad throughout the surgery. Mice were placed on a stereotaxic frame (David Kopf Instruments, USA), the skull was exposed, and the cranium was drilled with a micro motor handpiece drill (Foredom, USA: one or two holes for viral injection, two holes for screws with implantation, one or two holes for optic fibers, and one hole for a micro thermistor). The virus was injected unilaterally (right side) or bilaterally into the PBL (anteroposterior (AP), −1 mm from lambda; mediolateral (ML), ±1.5 mm; dorsoventral (DV), −3.5 mm, re-zero at the midline with the same AP). The viral injection was administered with a glass pipette (tips broken for an inner diameter of 20 μm) connected to the Nanoject III Programmable Nanoliter Injector (Drummond Scientific, USA) at a rate of 60 nL/min and retracted from the brain slowly after 5-10 min. For implantation, optic fibers were implanted above the injection site with the DV noted below, and the micro thermistor head was carefully lowered into the hole above the nasal cavity (AP +3.5 from the nasal fissure, ML 0.3). The implants were covered with superglue and dental cement for stabilization. Behavioral experiments were performed three weeks after viral injection and one week after the micro thermistor implantation unless otherwise noted.

For fiber photometry, *Oprm1*^*Cre*/+^ mice were unilaterally injected with 200 nL of either AAV-DIO-jGCaMP7f (1.5 E+14 GC/mL) or control AAV-DIO-eYFP (2.12E+12 GC/mL) into the PBL, and a stainless steel mono-fiber-optic cannula (400 μm diameter, 0.37 NA, Doric Lenses) was implanted 0.05 mm above the injection site.

For chemogenetics, 200 nL of either AAV-DIO-hM3Dq-mCherry (6.56E+11 GC/mL), AAV-DIO-hM4Di-mCherry (6.04E+11 GC/mL, Salk Institute Viral Vector Core), or control AAV-DIO-eYFP (2.12E+12 GC/mL) was injected bilaterally into the PBL of *Oprm1*^*Cre*/+^ (for hM3Dq) or *Oprm1^Cre/Cre^* (for hM4Di) mice.

For optogenetics, mice were bilaterally injected with 200 nL of AAV-DIO-ChR2-eYFP (1.8E+13 GC/mL) for photostimulation, 300 nL of AAV-DIO-ArchT-eYFP (1.26E+12 GC/mL, Salk Institute Viral Vector Core) for photoinhibition, or a corresponding amount of control AAV-DIO-eYFP (2.12E+12 GC/mL) into the PBL of *Oprm1*^*Cre*/+^ mice. A custom made mono-fiber-optic cannula (200 μm diameter, 0.22 NA) was implanted 0.3 mm above the injection site.

For monosynaptic rabies tracing of PBL^*Oprm1*^ neurons, 200 nL of AAV8-hSyn-FLEX-TVA-P2A-GFP-2A-oG (3.64E+13 GC/mL, Salk Institute Viral Vector Core) was injected unilaterally into the PBL. After three weeks, 200 nL of EnvA-ΔG-rabies-mCherry (3.95 E+08 GC/mL, Salk Institute Viral Vector Core) was unilaterally injected into the PBL. Mice were sacrificed seven days following injection.

For the investigation of PBL^*Oprm1*^ projections to the ventrolateral medulla, *Oprm1*^*Cre*/+^ mice were bilaterally injected with 200 nL of AAV2-phSyn1(S)-FLEX-tdTomato-T2A-SypEGFP-WPRE (2.23E+11 GC/mL, Salk Institute Viral Vector Core) into the PBL. Euthanasia and histology were performed three weeks following injection.

### Fiber photometry

A fiber photometry system (405 and 465 nm Fiber Photometry System, Doric Lenses Inc, Canada for Fig. 1; pyPhotometry, USA for Fig. 4) was used to record PBL^*Oprm1*^ neural activities. For the Doric system, GCaMP isosbestic fluorescence (405-nm excitation) and calcium-dependent fluorescence (465-nm excitation) were recorded at a sampling rate of 12 kHz, and data were analyzed with the Doric Neuroscience Studio software. For the pyPhotometry system, both channels were recorded with the 1-color time-division setting at 100 Hz, and data were analyzed with custom MATLAB scripts. F0 was calculated by a least mean squares fitting of the 405-nm channel in reference to the 465-nm channel, and ΔF/F was calculated as (F_465_-F_405_fitted_)/F_405_fitted_.

Cross-correlation analysis between the calcium signals and the respiration data was performed using the z-scored data and the MATLAB “xcorr” function using the normalized option, which allows autocorrelations at zero lag to equal 1. The area under the curve (AUC) was calculated with the MATLAB “trapz” function.

In awake behaving animals, concurrent measurements of PBL^*Oprm1*^ neural activity and respiration were recorded in the animal’s home cage for 20 min with a 55°C hot plate that was custom-built with a TC-720 thermoelectric temperature controller (TE Technology Inc., USA) for four min. In anesthetized animals, concurrent measurements of PBL^*Oprm1*^ neural activity and respiration were taken with the delivery of mechanical and thermal pain. For mechanical stimuli, 0 g and 300 g of mechanical pressure were delivered to pinch the tail tip using a dial tension gauge (ATG-300-1, Vetus Instruments, USA). For thermal stimuli, 25°C and 55°C heat were administered to the tail tip using a rod connected to a temperature controller (TA4-SNR+K, Mypin, China). Painful stimuli were delivered for 5 s after a stable 10-s baseline. AUC was calculated for 0-2.5 s before and 2.5-5 s after the mechanical or thermal stimulation, as well as during the 2-min segment before and after the start of the temperature change during the hot-plate assay.

### Convergent cross-mapping

State-space reconstruction models were generated using the framework of convergent cross mapping^3^, a nonlinear time series embedding method^4^ based on the Takens theorem and its generalized form^5^ that builds low-dimensional manifolds from time series and makes predictions across variables. Analysis and predictions were calculated using the R package rEDM 0.7.2 (https://cran.r-project.org/web/packages/rEDM/) for evaluation and the rEDM 0.7.4 (https://ha0ye.github.io/rEDM/) for model predictions in the RStudio environment. These packages were run on a dual Intel Xeon Gold 6148 Server with 384GB RAM or an Intel Core i9 2.4 GHz MacBook Pro with 32 GB RAM. Key parameters were determined individually by lagged coordinate embedding using the simplex function implementation in rEDM to optimize predictive skill as measured by the predicted over observed rho. Parameters include the delay tau, which gives the characteristic timescale of the time series, and the embedding dimensionality, which estimates the number of variables driving the system and approximates the real number of variables as given by the Whitney embedding theorem^6^ as minimally equal to the real number n of variables, but no more than two times n + 1 (*n* ≤ *E* ≤ 2*n* + 1). The choice of tau was also informed by minimizing mutual information^7^. This approximately corresponds to an autocorrelation of ~0.3, which was applied if it maximized predictive skill across datasets. To prevent data contamination, an exclusion radius was applied that was larger than the respiration rate smoothing window of five timesteps. Whenever the data allowed, an exclusion radius of E*tau was applied, unless the data was insufficient to apply this upper bound. In this case, the exclusion radius would be made just larger than tau. In this application CCM was used to generate multidimensional models from embeddings of jGCaMP7s activity that were able to predict respiratory rate using only jGCaMP7s activity.

### Chemogenetics

For behavioral testing, clozapine-N-oxide (CNO, Cayman Chemical, USA) was diluted in 0.9% saline to make a 1 mg/mL working solution. A final concentration of 5 mg/kg (for breathing-related tests) or 7.5 mg/kg (for all other behaviors without breathing) was injected intraperitoneally. Behavioral testing began 45-60 min after CNO injection. Animals that did not exhibit expression on one side were excluded from the analysis.

For medullary activation of PBL^*Oprm1*^ projections, clozapine (CLZ), a high-affinity agonist for Designer Receptors Exclusively Activated by Designer Drugs (DREADDs)^8^, was used to activate the PBL^*Oprm1*^ axon terminals in the ventrolateral medulla (VLM) respiratory center. Proper controls were included to ensure that CLZ selectively actives DREADD-expressing neurons. CLZ was first solubilized at pH 2.0 with HCl and then diluted. HPBCD [(2-Hydroxypropyl)-β-cyclodextrin] was added to solubilize clozapine with an HBPCD:CLZ molar ratio of 4:1. Solutions were titrated to neutral pH using NaOH and filter sterilized (0.22 μm) prior to intracranial administration^9^. CLZ (1 μg/μL) was injected bilaterally to the VLM with 100 nL/side using a glass pipette. A 50-nL aliquot of Cholera Toxin Subunit B-488 was injected using the same coordinates to mark the injection site. Data from 0 - 3 min before and 7 - 10 min after CLZ injection were used for quantification.

### Preparation of acute brain slices and electrophysiology

Mice were anesthetized with isoflurane, and the vascular system was perfused with ice-cold cutting solution (110.0 mM choline chloride, 25.0 mM NaHCO_3_, 1.25 mM NaH_2_PO_4_, 2.5 mM KCl, 0.5 mM CaCl_2_, 7.0 mM MgCl_2_, 25.0 mM glucose, 5.0 mM ascorbic acid and 3.0 mM pyruvic acid, bubbled with 95% O_2_ and 5% CO_2_). Mice were decapitated, and brains were quickly removed and chilled in ice-cold cutting solution. Coronal slices containing the PBL (250 μm) were cut using a VT 1200S Vibratome (Leica, Germany) and transferred to a storage chamber containing artificial cerebrospinal fluid (aCSF; 124 mM NaCl, 2.5 mM KCl, 26.2 mM NaHCO_3_, 1.2 mM NaH_2_PO_4_, 13 mM glucose, 2 mM MgSO_4_ and 2 mM CaCl_2_, at 32 °C, pH 7.4, bubbled with 95% O_2_ and 5% CO_2_). After recovery of at least 30 min, slices were transferred to room temperature (22 - 24 °C) for at least 60 min before use.

Slices were transferred into the recording chamber, perfused with aCSF at a flow rate of 2 mL/min. The temperature of the aCSF was controlled at 32 °C by a TC-324C Temperature controller (Warner Instruments, USA). PBL^*Oprm1*^ neurons were visualized with 490 nm epifluorescence illumination with a Scientifica SliceScope Pro (Scientifica, UK). Whole-cell and cell-attached patch-clamp electrophysiology was performed with Multiclamp 700B amplifiers (Molecular Devices, USA). Signals were digitized at 10 kHz with Digidata 1550B (Molecular Devices, USA).

For cell-attached, patch-clamp, aCSF was used as the internal solution. The spontaneous firing was measured before and after a 3 μM CNO perfusion. For whole-cell patch-clamp recordings, K^+^-containing internal solution was used (130 mM K-gluconate, 20 mM HEPES, 2 mM NaCl, 4 mM MgCl_2_, 0.25 mM EGTA, 4 mM Na-ATP, 0.4 mM Na-GTP, pH 7.2). Under current-clamp conditions, a 50-pA current was injected into the cell to induce action potential firing before and after a 3 μM CNO perfusion.

### Optogenetics

For ChR2 photostimulation and ArchT photoinhibition, a 470 nm collimated diode and a 589 nm diode-pumped solid-state (DPSS) laser systems (LaserGlow Tech., Canada) were used with a bilateral patch cord. The output power was 5 ± 1 mW as measured from the tip of the optic fiber. For investigating ChR2-induced breathing changes, 5-ms square pulses at 20 Hz were given for 10 s. The maximum breathing parameters 10 s before and 10 s after the stimulation onset were used for analysis. For investigating ArchT-induced breathing changes, a 30 s continuous-wave stimulation was delivered, and the average breathing parameters 10 s before and 10 s after the stimulation onset were used for analysis.

For the ArchT-stimulation combined with mechanical and thermal pain stimulation, 300 g of mechanical pressure and 55°C heat were administered to the mouse tail tip using a dial tension gauge (ATG-300-1, Vetus Instruments, USA) and a temperature controller (TA4-SNR+K, Mypin, China), respectively. Painful stimuli were given for 5 s after a stable 10-s baseline, and a subsequent 5-s photoinhibition was given for 3-5 s. For quantification purposes, three 3-s breathing episodes were analyzed: immediately before, immediately after the painful stimuli and during the 5-s photostimulation.

For the ArchT- and ChR2-stimulation combined with the real-time, place-preference (RTPP) or avoidance (RTPA) assay, a two-chamber arena (30 x 60 x 30 cm^3^) was used. A Gigabit Ethernet (GigE) USB camera (DFK 33GX236, Imagine Source, Germany; 25 frames/s, FPS), together with a video-tracking software (EthoVision XT 12, Noldus Information Technology Inc., USA), were used to track the animal and control the stimulation. After connecting to the patch cord, mice were placed in one side of the chamber for a 10 min baseline session and a subsequent 10 min test session, where a continuous wave of photoinhibition (ArchT), or 40 Hz photostimulation (ChR2) was given when mice go into one side of the chamber. The stimulation side was counterbalanced among animals. A 70% ethanol solution and deionized water were used for cleaning immediately after each test.

### Histology

Mice were euthanized with CO_2_ at a flow rate of 1.2 liters/min (LPM), perfused intracardially with ice-cold phosphate-buffered saline (PBS) and fixed with 4 % paraformaldehyde (PFA, 19210, Electron Microscopy Sciences, USA) in phosphate buffer (PB). The brain was extracted, post-fixed in 4% PFA overnight and dehydrated in 30 % sucrose in PBS until sliced. Frozen brains were cut into 50-μm coronal slices using a CM 1950 cryostat (Leica, Germany) and stored in PBS prior to mounting. The slices were mounted on Superfrost microscope slides (Fisher Scientific, USA) with DAPI Fluoromount-G mounting media (Southern Biotech, USA) for imaging.

### Immunohistochemistry

To investigate PBL^*Oprm1*^ projections onto preBötC neurons, anti-somatostatin (SST, a marker for preBötC) and anti-choline acetyltransferase (ChAT, a marker for motor neurons in the nearby nucleus ambiguous) staining was performed on both coronal (AP −7.3 mm) and sagittal (ML ±1.2 mm) sections. Mice were euthanized with CO_2_ at a flow rate of 1.2 LPM, then perfused intracardially with ice-cold PBS and then 4 % PFA in PB. The brain was extracted, post-fixed in 4 % PFA overnight and dehydrated in 30 % sucrose in PBS until sliced. Frozen brains were cut into 30-μm coronal slices with a CM 1950 cryostat and stored in PBS.

Sections were washed with PBST. After blocking with 3 % NDS for 1 h at room temperature and rinsing with PBST, slices were incubated with rabbit anti-SST14 (1:250, Peninsula Laboratories LLC, USA) and goat anti-ChAT (1:100, Sigma-Aldrich, USA) at 4 °C for 24 h. The next day, brain tissues were rinsed with PBST, then incubated in Cy3-conjugated Donkey Anti-Rabbit IgG and Alexa Fluor 647-conjugated Donkey Anti-Goat IgG (1:500 in 3 % NDS) for 90 min at room temperature. After washing with PBS, the slices were mounted on Superfrost microscope slides with DAPI Fluoromount-G mounting media for imaging.

### Imaging

All images were taken with the BZ-X710 all-in-one fluorescence microscope with the BZ-X viewer software under a 10X, 0.45 NA objective (Keyence, Japan), except for the Rabies Tracing experiments. Images for monosynaptic Rabies tracing were taken at the Salk Institute Waitt Advanced Biophotonics Core with the Olympus VS-120 Virtual Slide Scanning Microscope under an Olympus UPLSAPO 4X, 0.16 NA objective. For comparison, images were processed with the same gain, offset, and exposure time.

### Monosynaptic rabies tracing

Brain slice collection and region assignments were performed according to the Allen Brain Atlas. Every 50 μm section from AP + 2.62 mm to AP −7.455 mm was collected and imaged by an Olympus VS-120 Virtual Slide Scanning Microscope using the OlyVIA software using identical magnification and exposure time. Neurons with mCherry-positive cell bodies were counted manually. Brain slices located ±0.4 mm anterior and posterior to the starter region were excluded. The injection was unilateral (right side) and the quantification of cells was bilateral. The percentage of the total presynaptic input from a given brain region was calculated by dividing the number of presynaptic neurons by all neurons registered to the atlas.

### Elevated-plus maze

To measure anxiety, a custom-built plus-shaped maze with two 77-cm long opposite closed arms, two 77-cm long opposite open arms, and a central square of 7-cm sides was situated 70 cm above the ground. EthoVision XT 12 software and a GigE USB camera with 25 FPS were used to track the animal. The test mouse was introduced to the open arm adjacent to the center and was allowed to explore the maze for 10 min. The time spent and the entries in open arms during each session were calculated by the EthoVision XT 12 software. A 70% ethanol solution and deionized water were used for cleaning immediately after each test.

### Hot plate test

To measure thermal sensitivity, mice were placed inside a cylindrical, transparent Plexiglass chamber (D = 11 cm, H = 15 cm) on a hot plate (48°C or 55°C, PE34, IITC Life Science, USA). The latency of the painful response (hind paw shake, lick or jump) was manually recorded.

### Electronic von Frey test

To measure mechanical sensitivity, the Dynamic Plantar Aesthesiometer (37450, Ugo Basile, Italy) was used with the maximum force set to reach 50 g after 20 s. Mice were habituated inside a Plexiglass chamber (10 x 10 x 13 cm) on a metal mesh floor for >2 h until they were awake but largely immobile. The metal rod was placed underneath the left hind paw of the mice. The system automatically measures the latency and force delivered upon mice hind paw withdrawal. This measurement was taken 5 times with >1 min interval. The averaged value was used for further analysis.

### Formalin test

To measure affective pain responses, 4% formalin (1.6% PFA) was injected subcutaneously in the left hind paw of an awake mouse. This produced a biphasic pain response over a 1-h test period. Mice were placed in a Plexiglass chamber (10 x 10 x 13 cm) with a mirror placed behind it. Pain responses, such as licks, twitches, raising or shaking of the injected paw, were scored. The percentage of time during the acute (0-5 min following injection) and the inflammatory phase (20-35 min following injection) were used for quantification.

### Contextual fear conditioning

The fear-conditioning chamber (ENV-007CT, MED Associates INC, USA) is an arena (26 x 30 x 33 cm) with two Plexiglass walls, two metal walls, and a metal grid floor to deliver electrical shocks (ENV-005, MED Associates INC). The chamber was connected to a standalone aversive stimulator (ENV-414S, MED Associates INC) and enclosed in a light- and sound-attenuating cubicle (ENV-018MD, MED Associates INC). EthoVision XT 12 software and a GigE USB camera with 25 FPS were used to track the animal. A 70% ethanol solution and deionized water were used for cleaning immediately after each test. On day 1 and day 2, mice underwent two 6-min habituation sessions in the chamber. On day 3, mice were introduced to the chamber 45-60 min after CNO injection and subsequently received five shocks (2 s, 0.2 mA) with uneven intervals during a 7-min trial. After 24 h, mice were reintroduced to the chamber for a 2-min context-dependent retrieval test. The percentage of time freezing and the total distance moved during each session were calculated by the EthoVision XT 12 software. Freezing behavior was defined as the period during which the velocity of the mouse was less than 1.75 cm/s for at least 3 s. Automatic scoring was validated with manual scoring.

### Statistical analysis

All data are shown as mean ± SEM and analyzed using either a student’s t-test, one-way ANOVA with Tukey’s *post hoc* comparison, or two-way ANOVA with Bonferroni’s *post hoc* comparison. All the statistical analyses were performed with Prism 6 (GraphPad Software Inc., USA). ns, not significant; *, *p* < 0.05; **, *p* < 0.01; ***, *p* < 0.001; ****, *p* < 0.0001.

### Data availability

The data that support the findings of this study are available from the corresponding author upon reasonable request.

---

**Figure S1.**
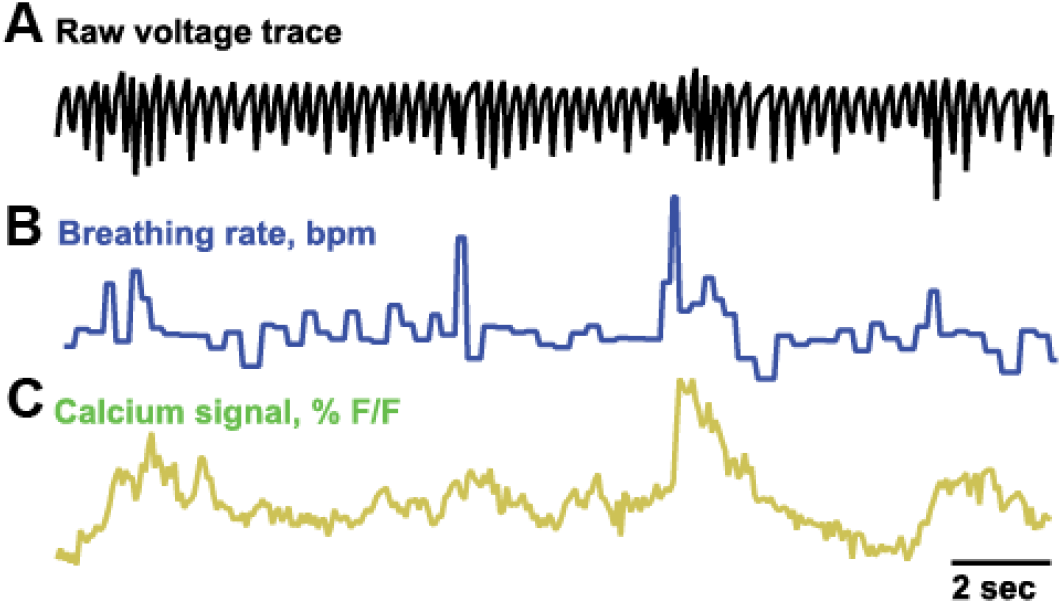
**A-C**, Example (**A**) raw voltage traces from the micro thermistor, (**B**) breathing rate and (**C**) calcium signals of PBL^*Oprm1*^ neurons for a 25 sec period.

**Figure S2.**
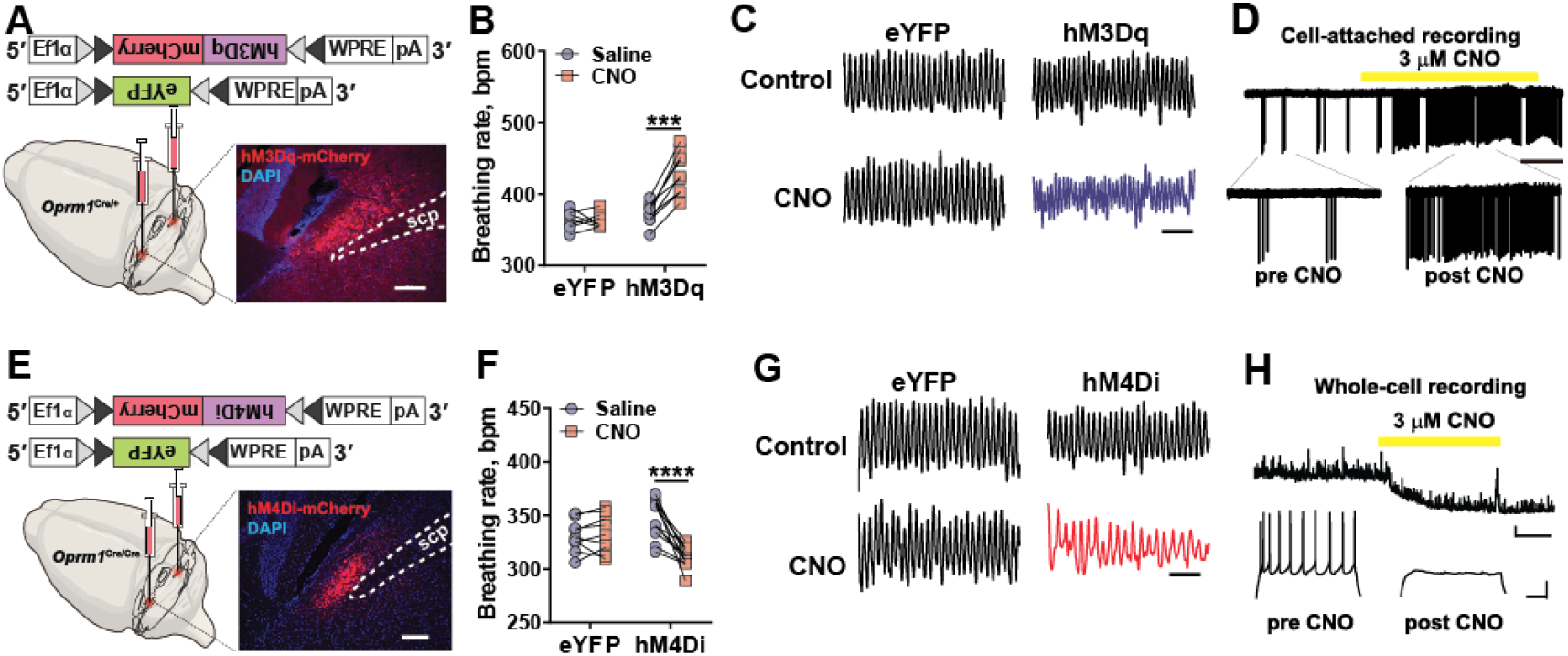
**A**, Stereotaxic injection of AAV-DIO-hM3Dq-mCherry and AAV-DIO-eYFP into the PBL of the *Oprm1*^Cre/+^ mice. **B**, Whole body plethysmography (WBP) showed that injection of CNO (5 mg/kg) significantly increased breathing rates in hM3Dq-expressing awake mice (n = 9), whereas eYFP-expressing mice (n = 7) displayed no significant changes in breathing rates by CNO. Two-way ANOVA analysis with Bonferroni’s multiple comparison post-hoc test, ***, *p* < 0.001. **C**, Example breathing traces after systemic injection of control (0.9% saline) and CNO in eYFP- and hM3Dq-expressing mice. Scale bar, 1 sec. **D**, CNO-induced activation of hM3Dq-expressing PBL^*Oprm1*^ neurons in *ex vivo* cell-attached recording. Scale bar, 20 sec. **E**, Stereotaxic injection of AAV-DIO-hM4Di-mCherry and AAV-DIO-eYFP into the PBL of the *Oprm1*^Cre/Cre^ mice. **F**, WBP showed that injection of CNO (5 mg/kg) significantly decreased breathing rates in hM4Di-expressing awake mice (n = 12), whereas eYFP-expressing mice (n = 9) displayed no significant changes in breathing rates after CNO. Two-way ANOVA analysis with Bonferroni’s multiple comparison post-hoc test, ****, *p* < 0.0001. **G**, Example breathing traces after systemic injection of control (0.9% saline) and CNO in eYFP- and hM4Di-expressing mice. Scale bar, 1 sec. **H**, CNO-induced inhibition of hM4Di-expressing PBL^*Oprm1*^ neurons in *ex vivo* whole-cell patch-clamp recording. Scale bar, 2 mV / 20 sec.

**Figure S3.**
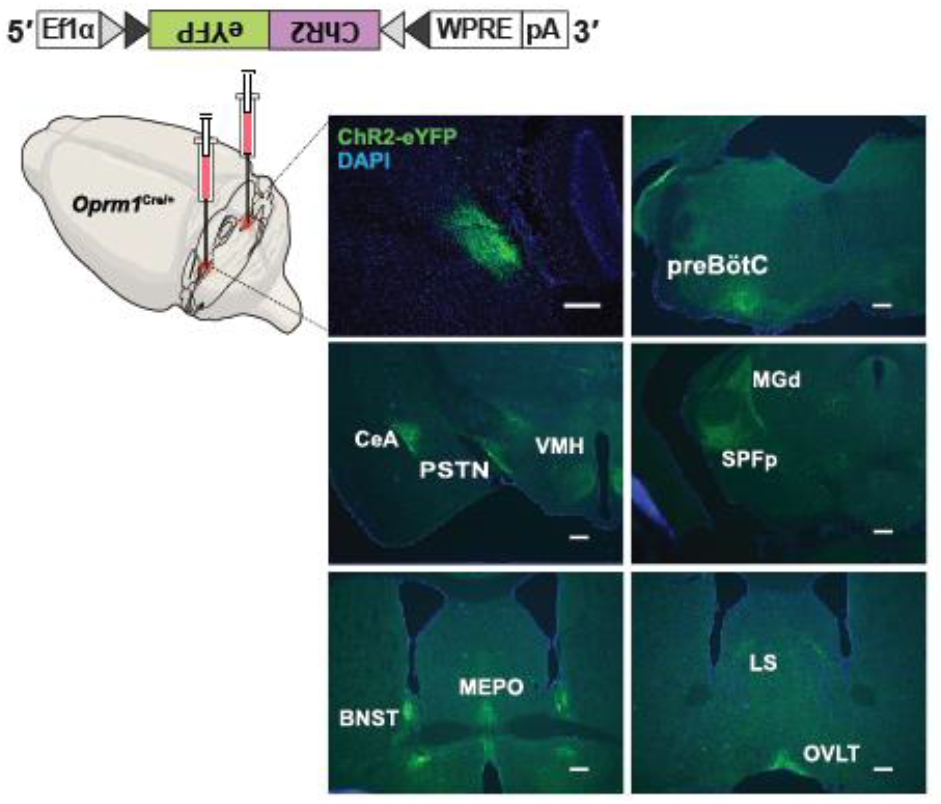
Output mapping of the PBL^*Oprm1*^ neurons by stereotaxic injection of AAV-DIO-ChR2-eYFP into the PBL of *Oprm1*^*Cre*/+^ mice (top left panel) and visualization of the axon terminals in example brain regions. Abbreviations: preBötC: pre-Bötzinger complex; CeA: central amygdala; PSTN: parasubthalamic nucleus; VMH: ventromedial hypothalamus; MGd: medial geniculate complex, dorsal part; SPFp: subparafascicular nucleus, parvicellular part; BNST: bed nuclei of the stria terminalis; MEPO: median preoptic nucleus; LS: lateral septal nucleus; OVLT: organum vasculosum of the laminae terminalis. Scale bar, 200 μm.

**Figure S4.**
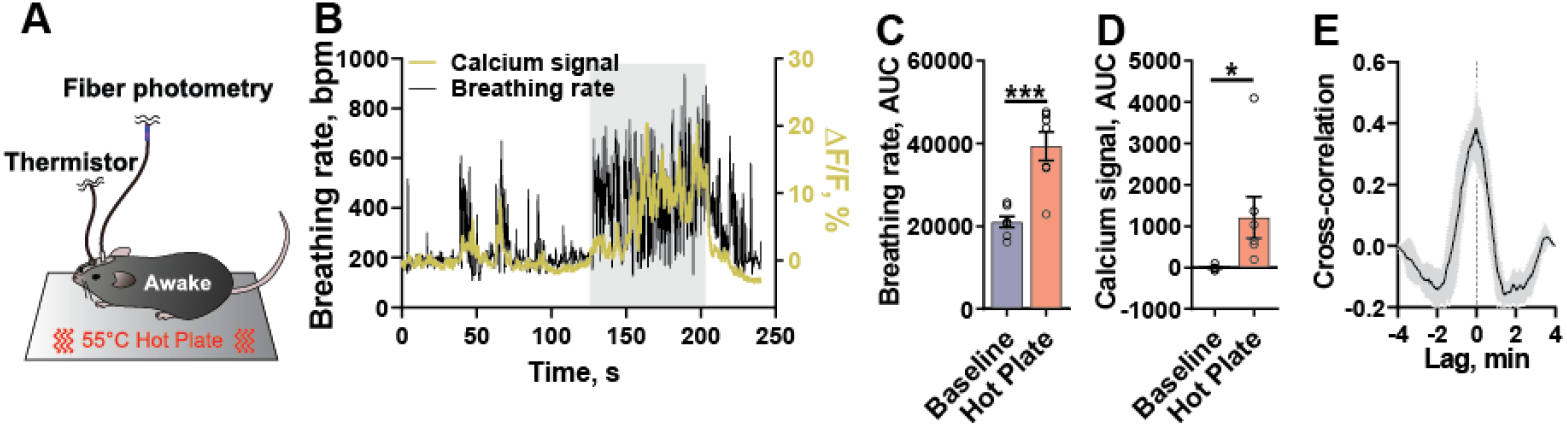
**A**, Simultaneous monitoring of PBL^*Oprm1*^ neuronal activity and breathing rate when freely moving mice were stimulated with thermal-pain signal (55°C hot plate). **B**, Traces of breathing rate and calcium signal in a test mouse, gray area, hot temperature applied. **C**, AUC analysis of the breathing rate (n = 7), unpaired t-test, ***, *p* < 0.001. **D**, AUC analysis of the calcium signals (n = 7), unpaired t-test, *, *p* < 0.05. **E**, Cross-correlation analysis of breathing rates and calcium signals (n = 7).

**Figure S5.**
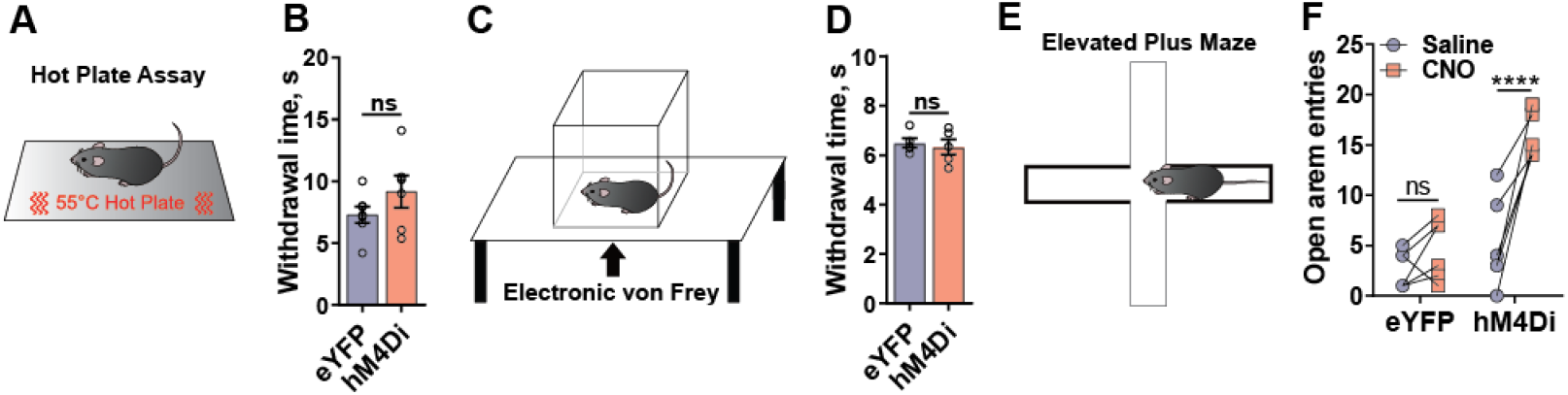
**A-B**, Hot-plate test for thermal sensitivity in mice expressing eYFP or hM4Di in PBL^*Oprm1*^ neurons (n = 7). ns, not significant, unpaired t-test. **C-D**, Electronic Von Frey test for mechanical sensitivity in mice expressing eYFP or hM4Di in PBL^*Oprm1*^ neurons (n = 5). ns, not significant, unpaired t-test. **E**, Elevated plus maze (EPM) test with mice expressing eYFP or hM4Di in the PBL^*Oprm1*^ neurons. **F**, Mice expressing hM4Di displayed more open arm entries when injected with 7.5 mg/kg CNO, as compared with saline injection, whereas the eYFP-expressing control group showed no significant increase by CNO, or Saline injection (n = 7). Two-way ANOVA analysis with Bonferroni’s multiple comparison post-hoc test. ns, not significant; ****, *p* < 0.0001.

